# Inhibition of CHK2 dependent DNA damage response suppresses Zika virus infection through a STING dependent mechanism

**DOI:** 10.1101/2022.10.03.510601

**Authors:** Huailiang Ma, Chunfeng Li, Yanli Wang, Erika V. Valore, Meera Trisal, Harold Sai-yin Hui, Paul Hakimpour, James R. Bowen, Tak W Mak, Mehul S. Suthar, Bali Pulendran

## Abstract

Zika virus (ZIKV) is a mosquito-borne flavivirus that causes neurological disorders and microcephaly. Recent research has shown that ZIKV causes cell cycle arrest and DNA damage response in neuronal progenitor cells that potentially leads to congenital neurodevelopmental defects. However, the specific role of regulators that control DNA damage response to ZIKV infection and the related mechanisms remain largely unknown. Here through both *in vitro* and *in vivo* studies, we observe that ZIKV induces DNA damage response in both human cell line and mouse embryo. When CHK2 dependent DNA damage response pathway is inhibited by a small molecule inhibitor or genetic deletion, ZIKV production is reduced and the embryonic developmental defects are rescued. Furthermore, we demonstrate that in ZIKV infected *Chk2-/-* mice, the reduced viral load correlates with elevated antiviral innate immune response which is found to be dependent on the STING signaling pathway. Collectively, our study reveals a specific role of CHK2 for ZIKV infection and pathogenesis, demonstrating a mechanism that inhibition of the CHK2 axis suppresses ZIKV infection through the STING-dependent antiviral pathway, thus providing a new therapeutic strategy against ZIKV. These findings also suggest the intriguing possibility that ZIKV evolved to orchestrate DNA damage response factors to create a beneficial environment for its infection in the host cells.

## Introduction

Zika virus (ZIKV) is the causative agent of Zika fever, which is characterized by rashes, conjunctivitis and headache ^1^. ZIKV infection during pregnancy causes intrauterine growth restriction and microcephaly as well as fetal brain abnormalities, but the underlying mechanisms are not fully understood. Several studies have demonstrated that ZIKV can infect macrophages and replicate efficiently in the placenta ^2, 3^. After successfully bypassing this natural barrier, ZIKV preferentially propagates in the fetal neural progenitors and causes cell cycle arrest ^4, 5^. Ultimately, this may impair the differentiation and growth of fetal neurons, contributing to the brain developmental defects seen in the newborns ^5, 6, 7^. It is also possible that maternal ZIKV exposure may directly damage the placental tissue where trophoblast hyperplasia in the labyrinth and trophoblast giant cell necrosis in the junctional zone have been reported ^8, 9^. The resulted lack of nutrition supply in placenta may indirectly restrict the cell cycle progression of neural progenitors ^9^. Given the unique biology of ZIKV in manipulating cell cycle arrest compared to other flaviviruses in the same family ^4, 5^, there is much interest in understanding the specific cellular and molecular mechanisms involved.

Global gene expression analysis of ZIKV-infected human neuronal precursor cells (hNPCs) and murine fetal brain tissues shows that some cell cycle related gene networks are notably dysregulated ^4, 10^. Of interest, p53, a key regulator in mediating genotoxic stress and cell cycle arrest, is found to be activated upon ZIKV infection ^11^. Several other genotoxic stress related proteins, such as ataxia-telangiectasia mutated (ATM), ATR (ATM and Rad3-related) and H2AX, are significantly phosphorylated in the ZIKV infected cells ^12^. Since ATM, ATR and γH2AX are the major regulators of the host DNA damage response (DDR) pathway, which in turn controls cell cycle progression, it is speculated that DDR regulators would have a potential role in regulating ZIKV infection and pathogenesis. However, the related functional data are not yet reported.

The cellular DDR pathway is composed of complex signaling networks that safeguard the cell in maintaining genomic DNA integrity during replication and when the cells are under endogenous and exogenous threats ^13, 14^. When DNA damage occurs, the transducer kinase ATM is recruited and activated at the site of double stranded DNA breaks (DSBs) and phosphorylates several downstream effector proteins, including the cell cycle checkpoint proteins CHK2, p53 and histone H2AX ^15^. For DNA single-strand breaks (SSBs) at the stalled replication forks, another transducer kinase ATR is primarily activated to phosphorylate its respective effector proteins CHK1 and p53 ^14^. These two major DDR pathways work in conjunction and are tightly regulated to control cell cycle at three checkpoints: G1/S, intra-S and G2/M phases, thus providing cells sufficient time to repair the damaged DNA before reentering the cell cycle ^14^.

Recently, Hammack et al have shown that ZIKV but not dengue virus (DENV) activates ATM-CHK2-γH2AX DDR and impedes cell cycle progression through S phase in hNPCs. Using inhibitors to arrest cell cycle at S-phase leads to an increase in ZIKV replication ^16^. However, the direct role of ATM or CHK2 in ZIKV infection is not explored, nor to mention the contribution of these DDR regulators to ZIKV pathogenesis *in vivo*. Here, we find that γ-H2AX, a hallmark for DNA DSBs is chiefly observed at the placenta and the developing fetus upon ZIKV infection. Blockade of CHK2 dependent DDR with small molecule inhibitor results in a substantial reduction of ZIKV production, and a concomitant rescue of fetal developmental defects. Importantly, in *Chk2*^*-/-*^ mice, ZIKV infection results in reduced virus production in different regions of fetal brain, further supporting the direct role of CHK2 in promoting ZIKV infection and pathogenesis. Furthermore, we reveal that there is an elevated antiviral innate immune response in *Chk2*^*-/-*^ mice upon ZIKV infection, which is found to be dependent on the STING signaling pathway through the studies from *Chk2-/-Sting-/-* mice.

## Results

### ZIKV infection triggers DDR both *in vitro* and *in vivo*

To confirm if ZIKV could uniquely induce DDR in regular lab-cultured human cell line rather than the sophisticated cultured hNPCs or embryonic stem cells^16^, we compared ZIKV infection in parallel with DENV-2 and yellow fever virus 17 strain (YF-17D) in Hela cells. After 36 hours infection at MOI of 0.1, the cells were stained with a pan-flavivirus antibody (anti-E) and the confocal images indicated that the percentage of viral-E antigen positive cells between ZIKV-PR and DENV-2 was comparable, while YF-17D exhibited a higher infection rate (about 2.1-fold) (**Fig**. 1A and **Fig**. 1B). In consistent with the previous finding ^17^, YF-17D infection could also induce stress granule, marked by G3BP1, in Hela cells (**Fig**. 1A). Interestingly, by western blot for studying the changes of key regulators in DDR pathway, we found that only ZIKV-PR could enhance the phosphorylation of ATM, CHK2 and γH2AX (**Fig**. 1C). Our data from Hela cells also reveals a similar effect of ZIKV to arrest cell cycle at S phase and modulate multiple cell cycle regulators ^16^ (**Fig**. S1). To further explore if ZIKV infection would induce DNA damage response *in vivo*, we utilized a pregnant mouse model developed by Miner et al. ^9^ (**Fig**. 1D). After virus infection at E8.5 on the *Ifnar1*^*-/-*^ dams that have mated with the wildtype BL6 mice, approximately half of the embryos (*Ifnar1*^*+/-*^) were found to be resorbed at day 7 post infection (**Fig**. 1E). From those un-resorbed embryos, we observed that the phosphorylation level of γH2AX was notably elevated at both placenta and fetus compared to those from mock infected groups (**Fig**. 1F-G). Together, these results demonstrate that ZIKV infection could induce host DDR not only *in vitro*, but also *in vivo*.

**Figure 1.**
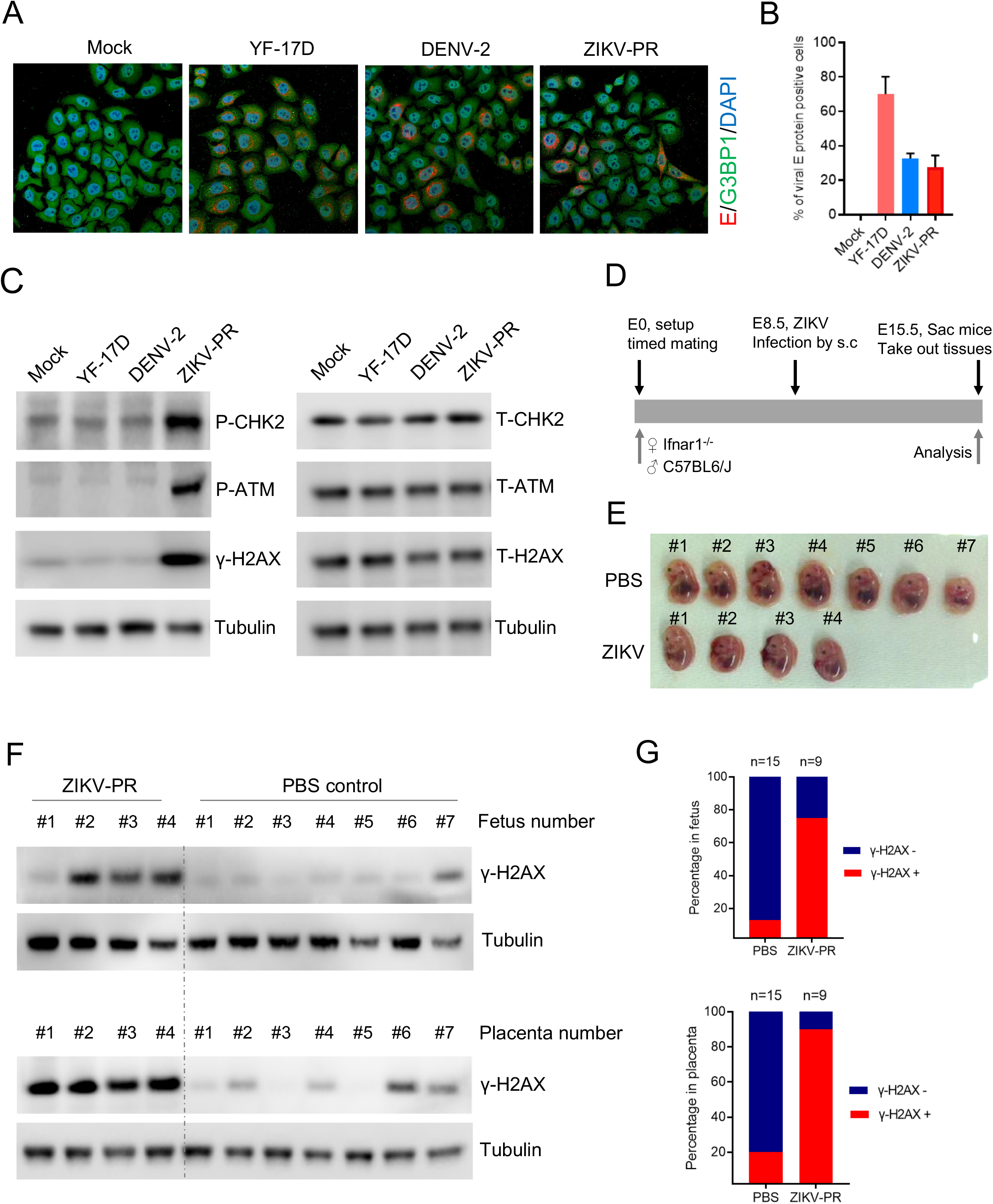
ZIKV induces host cell DNA damage response both *in vitro* and *in vivo*. **(A-B)** Hela cells were infected with ZIKV-PR, DENV-2 and YF-17D viruses at MOI of 0.1 for 36 hours. Then the cells were fixed, and viral E protein was stained by a pan-flavivirus antibody. The percentage of viral E protein positive cells from 3 independent experiments was quantified and plotted. G3BP1 is a marker of stress granule. **(C)** Hela cell lysates from the same repeated experiment in Fig. 1A were subjected to western blot analysis of the key regulators of DNA damage response signaling pathways. **(D)** Schematic of ZIKV infection in a pregnant mice model. The timed mating was setup for female *Ifnar1*^*-/-*^ and male WT mice, the pregnant dam were infected by ZIKV at E8.5 or inoculated with PBS as control. **(E)** After 7 days post infection, the dams were sacrificed, and all the intact embryos (*Ifnar1*^*+/-*^) were collected. **(F)** From the embryos, the fetuses and the corresponding placentas were homogenized in RIPA cell lysis buffer respectively, then subjected to western blot analysis of cell DNA damage response. **(G)** The proportion of fetus or placenta with DNA damage positivity was graphed. One representative data set from two or three independent experiments was shown here.

### Blockade of CHK2 pathway with small molecule inhibitor suppresses ZIKV infection both *in vitro* and *in vivo*

To investigate the direct role of DDR regulator to ZIKV infection and pathogenesis, we assessed two available small molecule inhibitors that target CHK2 and ATM (**Fig**. 2 and **Fig**. S2). With increasing doses of a CHK2 inhibitor, the phosphorylation level of CHK2, ATM and γH2AX was suppressed substantially in ZIKV infected Hela cells (**Fig**. 2A-B). Surprisingly, CHK2 inhibitor could also efficiently reduce ZIKV infected cells from 90% to approximately only 10% at a dose of 50µM (**Fig**. 2C-D). Furthermore, the infectious titer of ZIKV in the supernatant was decreased more than 3-logs after CHK2 inhibitor treatment (**Fig**. 2E). In contrast, the CHK2 inhibitor exhibited no antiviral effects for DENV-2 or YF-17D viruses (**Fig**. S2). Therefore, these data imply that targeting CHK2 dependent DDR pathway could uniquely inhibit ZIKV infection.

**Figure 2.**
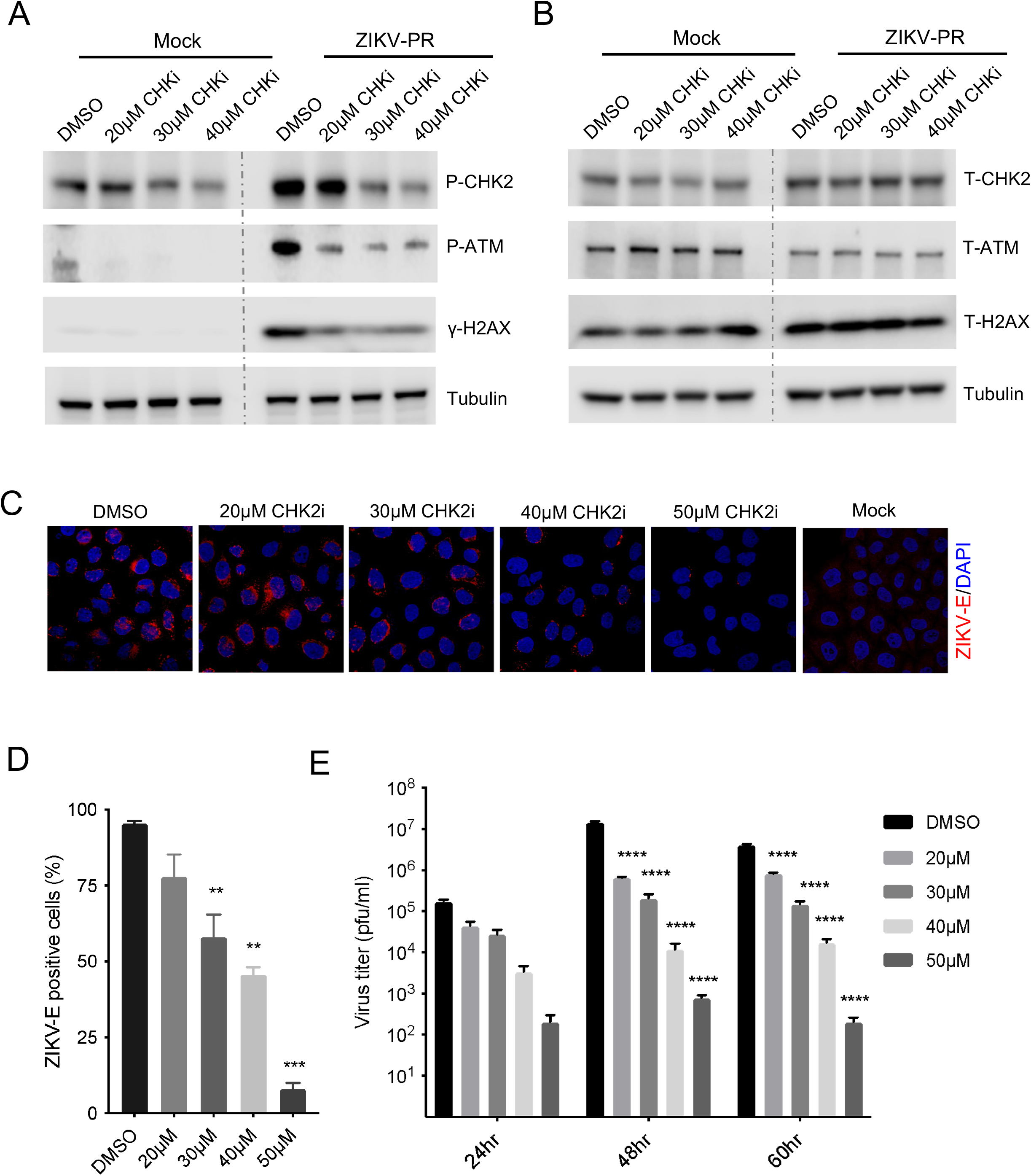
Pharmacological inhibition of CHK2 blocks ZIKV induced DNA damage response and dampens virus infection *in vitro*. **(A-B)** Hela cells were infected with ZIKV at MOI of 0.1 for 2 hours. After washing out of the unbound viruses, fresh culture medium with different doses of CHK2 inhibitor was added. The cell lysates were collected at 48 hours post infection, and the key regulators of cell DNA damage response were analyzed by western blot. **(C)** Hela cells from the same repeated experiment in Fig. 2A were fixed, and viral E protein was stained. **(D)** The percentage of viral E protein positive cells was quantified and plotted. **(E)** Hela cells were infected with ZIKV at MOI of 0.1, followed by CHK2 inhibitor treatment at different doses, the supernatants were collected for virus quantification by plaque assay at different time points. Error bars are mean ± SEM, average of three biological replicates from three independent experiments, * p<0.05, ** p<0.01, *** p<0.001, and **** p<0.0001.

Next, we determined if the observed anti-ZIKV effect of CHK2 inhibitor *in vitro* could be translated into any anti-viral effect *in vivo*. The pregnant *Ifnar1*^-/-^ dams that have mated with wildtype C57BL6/J males were infected with ZIKV at E8.5 by footpad subcutaneous injection as developed by Miner et al. ^9^ (**Fig**. 3A). CHK2 inhibitor was given daily by intraperitoneal injection starting on the same day as viral infection. CHK2 inhibitor treated dams gained more weight than the dams from the vehicle treated control group (**Fig**. 3B). At day E15.5 all dams were sacrificed, and we observed that from the ZIKV infected dams, CHK2 inhibitor treated groups had 25% more intact embryos (*Ifnar1*^+/-^) than the vehicle treated group (**Fig**. 3C-D). We also evaluated the effect of CHK2 inhibitor on the embryo development in absence of ZIKV infection and found both the vehicle and CHK2 inhibitor treated dams had the similar intact/resorbed embryo profile (**Fig**. 3C-D). Furthermore, from the ZIKV infected group, the average weight of the embryos from CHK2 inhibitor treated dams was significantly higher than that from the vehicle treated dams (**Fig**. 3E). Importantly, the viral loads in the placenta were significantly reduced in the CHK2 inhibitor treated group (**Fig**. 3F). Taken together, these data demonstrate that the CHK2 inhibitor have a similar anti-ZIKV effect *in vivo*.

**Figure 3.**
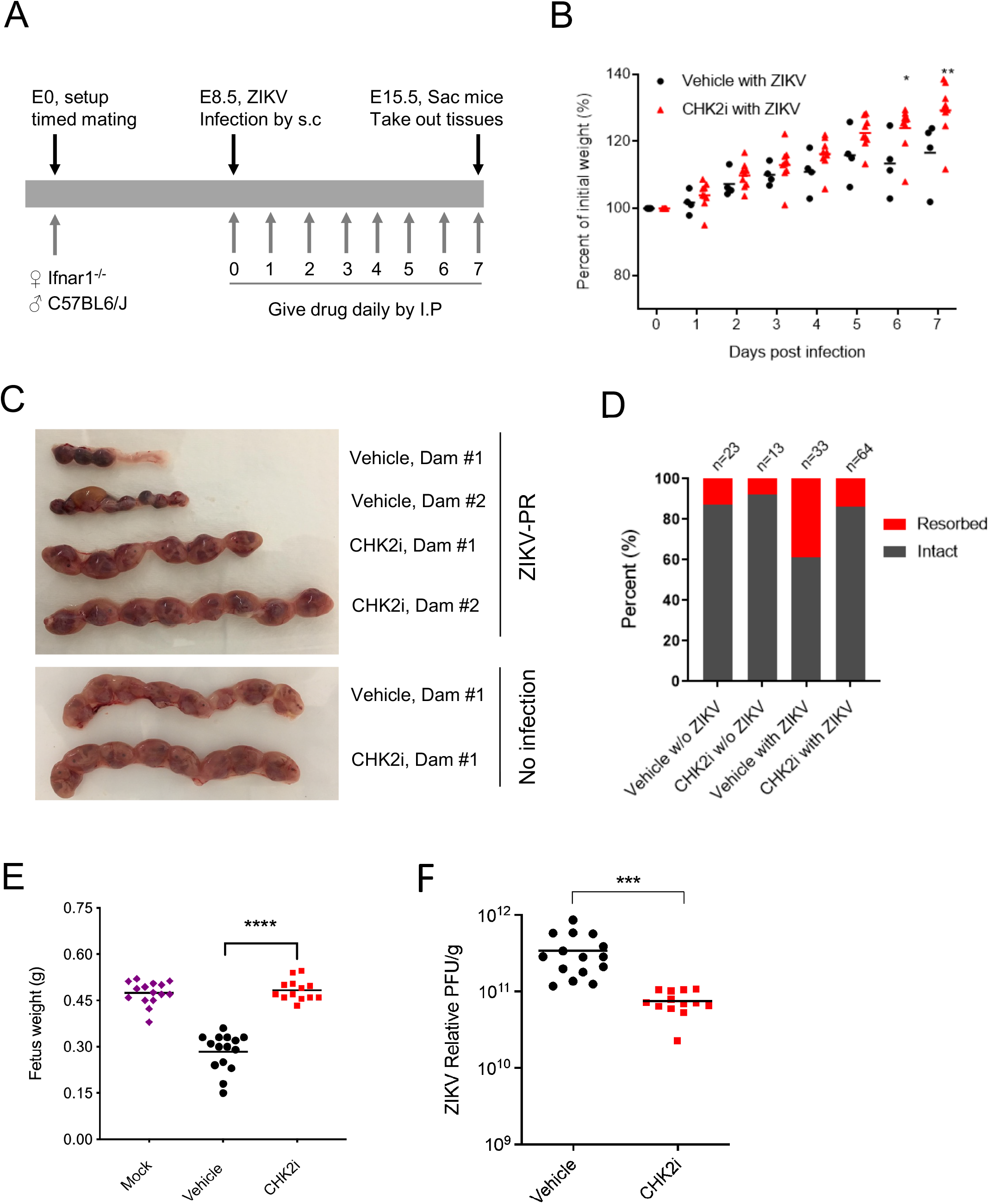
Pharmacological inhibition of CHK2 rescues ZIKV induced embryos resorption and reduces viral loads at placenta. **(A)** Schematic of ZIKV infection and CHK2 inhibitor treatment in a pregnant mice model. **(B)** The weight of ZIKV infected pregnant dam was recorded daily, and the changes of in body weight were normalized and plotted. **(C)** The embryos from one experiment were shown. **(D)** The percentage of resorbed embryos and **(E)** the weight of fetus from three independent experiments were plotted. **(F)** The viral loads at placenta were analyzed by qRT-PCR from three independent experiments, * p<0.05, *** p<0.001, and **** p<0.0001, One-way ANOVA (E) or student *t* test (F).

### Deletion of *Chk2* rescues mice from ZIKV induced neuropathogenesis

We further investigated the role of CHK2 in ZIKV-induced neuropathogenesis in a neonatal mouse model via intracerebral infection (**Fig**. 4A). The development of *Chk2*^*-/-*^ adult mice and neonatal mice is normal as described before ^18^. The data showed that ZIKV viral load in the brain of *Chk2*^*-/-*^ mice was significantly decreased over 1-log compared to those from wildtype mice (**Fig**. 4B). Additionally, a significant reduction of viral E protein positive cells was observed in the *Chk2*^*-/-*^ group as compared to wildtype control group (**Fig**. 4C). Consistently, higher survival rate was observed in ZIKV infected *Chk2*^*-/-*^ mice (**Fig**. 4D). ZIKV induced brain developmental defects observed in the wildtype neonatal mice was also found to be rescued in the *Chk2*^*-/-*^ mice (**Fig**. 4E-F).

**Figure 4.**
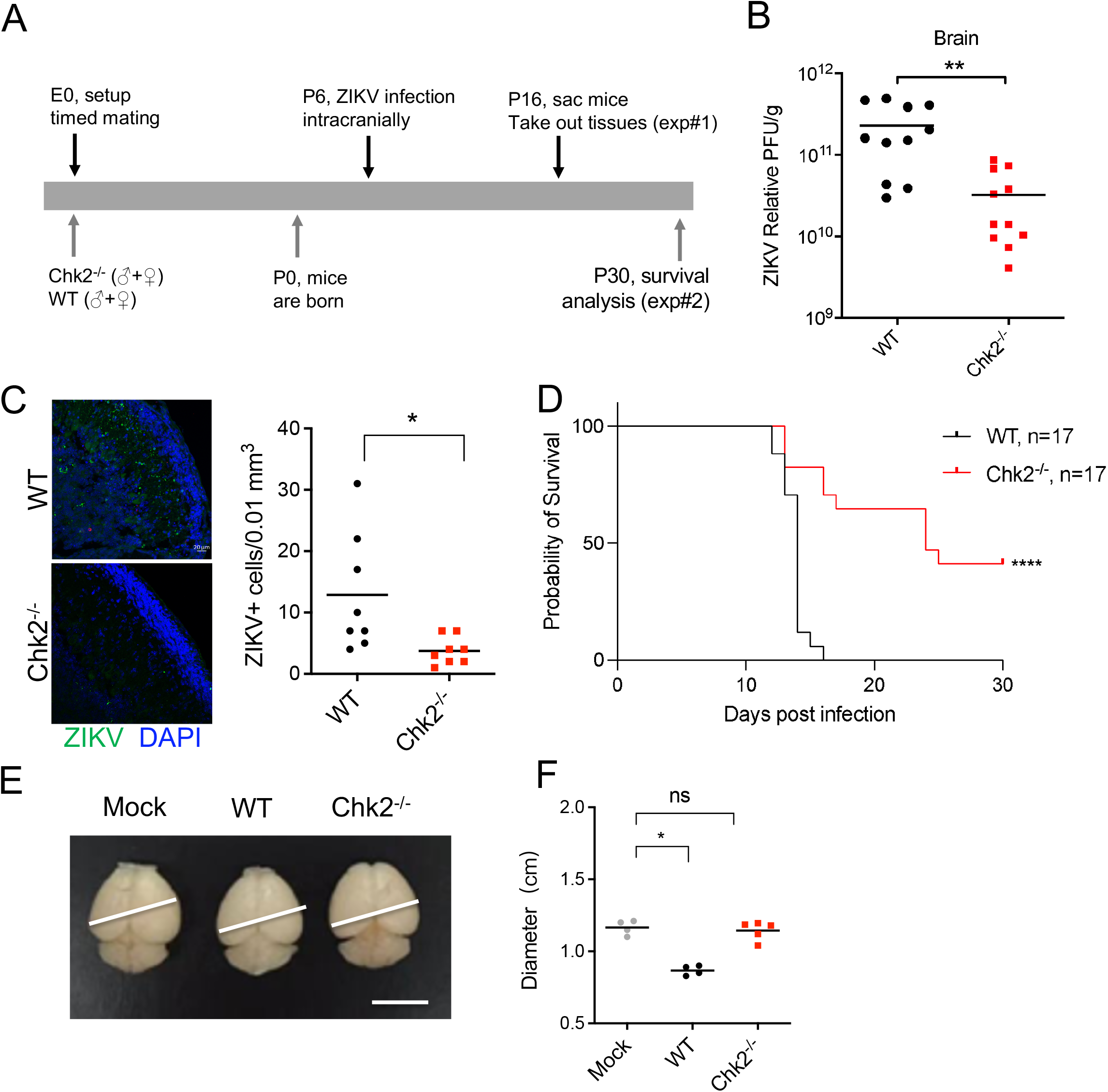
CHK2 is essential for ZIKV-induced neuropathogenesis in neonatal mice. **(A)** Schematic of ZIKV infection in a neonatal mice model. **(B)** Neonatal *Chk2*^-/-^ mice or littermate controls were infected with ZIKV at P6 by intracranial injection (ZIKV-PR strain, 100 PFU/mice). Viral loads in the brain of neonatal mice were detected by qRT-PCR at P16 from two independent experiments. **p<0.01, student *t* test followed by Mann-Whitney test. **(C)** Viral antigen in neonatal brain tissue section were detected by immunofluorescence assay. E protein is in green, Caspase3 is in red. Bar=20 µm. **(D)** The survival curve from the infected neonatal mice was plotted from 3 litters. WT (n=17), *Chk2*^*-/-*^ (n=17), ***, p<0.001, Log-rank (Mantel-Cox) test. **(E)** The size of brain from WT, *Chk2*^-*/-*^mice at P16. Non-infected WT mice brain was shown as control (Mock). Bar=1cm. **(F)** Diameter of brain from Mock, WT, and *Chk2*^*-/-*^ neonatal mice infected by ZIKV as shown in (A). *** p<0.001, One-way ANOVA (F) test. One representative data set from at least two independent experiments was shown in (**C, E-F**).

Li *et al*. showed that ZIKV can infect adult mice by targeting adult neuronal cells, especially the NPCs ^19^. Thus, we used the adult mouse intracranial infection model to study the role of CHK2 for ZIKV induced neuropathogenesis (**Fig**. 5A). We observed that ZIKV infection in *Chk2*^*-/-*^ mice was suppressed in different parts of the adult brain compared to the littermate control mice (**Fig**. 5B). Consistently, the clinical disease outcome and the survival profile of the *Chk2*^*-/-*^ mice were improved as well (**Fig**. 5C-D). Interestingly, there was no significant difference between WT and *Chk2*^*-/-*^ mice with DENV-2 and YF-17D infections, in agreement with the *in vitro* results (**Fig**. S3). Taken together, these data further confirm the direct role of CHK2 dependent DDR pathway in controlling ZIKV infection, vertical transmission and neuropathogenesis, but not in controlling DENV or YF-17D infection.

**Figure 5.**
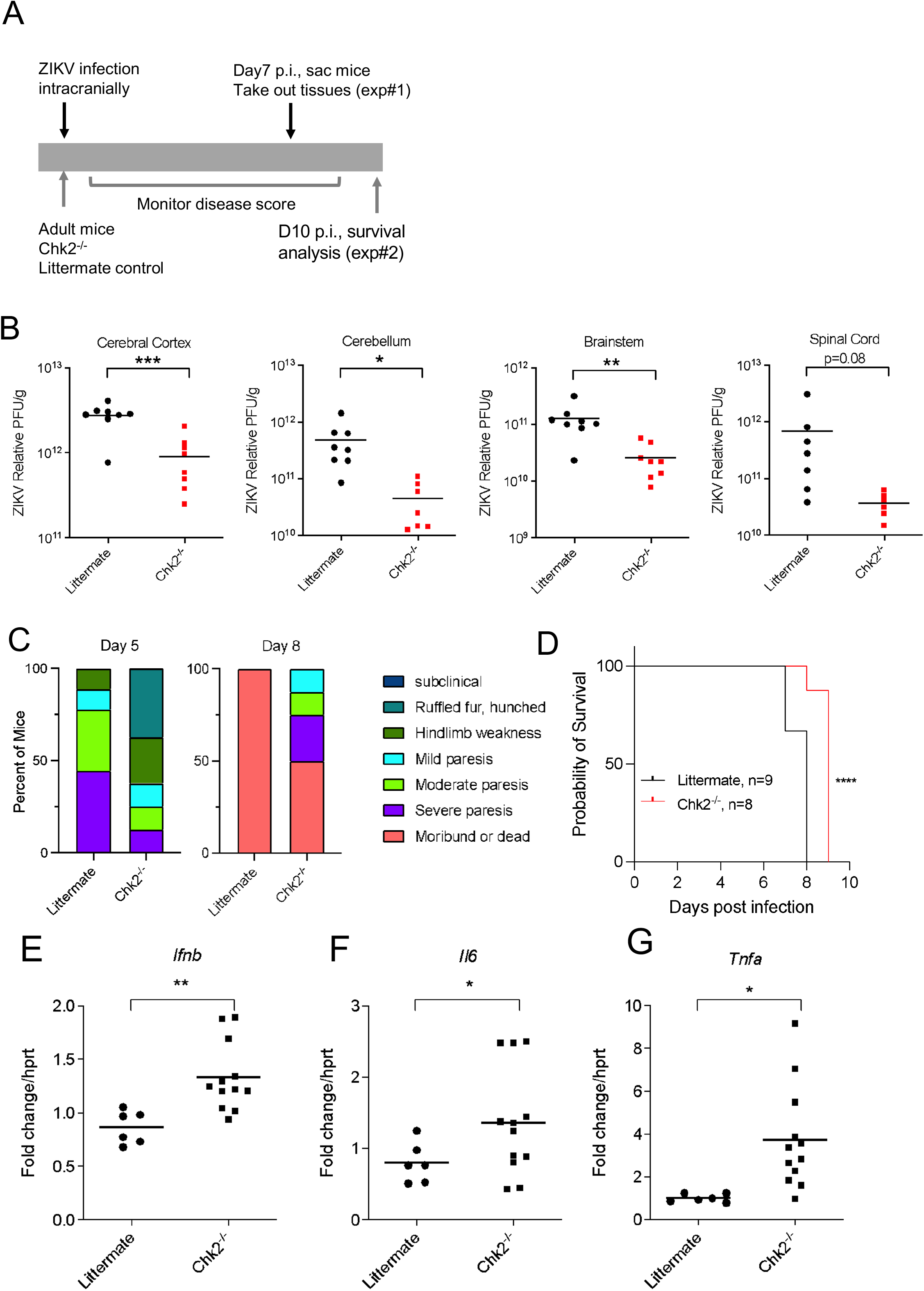
CHK2 is essential for ZIKV infection and pathogenesis in adult mice. **(A)** Schematic of ZIKV infection in the brain of adult mice. **(B)** Adult *Chk2*^-/-^ mice or littermate controls (6 weeks old) were infected with ZIKV (MR766 strain, 10^4^ PFU/mice) by intracranial injection. At 7 days post infection, viral loads in the cerebral cortex, cerebellum, brainstem, and spinal cord were detected by qRT-PCR. WT (n=8), *Chk2*^-/-^ (n=8), were collected from 3-4 litters respectively. **p<0.01, student t test followed by Mann-Whitney test. **(C)** The disease condition of each mouse was monitored and plotted according to different clinical symptoms at day 5 and day 8 post infection. **(D)** The Survival curve of the infected mice was plotted. WT (n=9); *Chk2*^*-/-*^ (n=8), **** p<0.0001, Log-Rank (Mantel-Cox) test. (**E-G**) *Ifnb, Il6*, and *Tnfa* gene expression in WT and *Chk2*^*-/-*^ mice brain infected by ZIKV (MR766 strain, 10^4^ PFU/mice) at 7 days post infection. *p<0.05, **p<0.01, student *t* test. Data was combined from two repeats.

### Inhibition of Chk2 axis promotes STING-dependent antiviral effect to suppress ZIKV infection and pathogenesis

Finally, we determined the mechanism of how pharmacological or genetic blockade of CHK2 axis reduced the viral load and rescued ZIKV induced pathogenesis. Thus, we first assessed interferon beta production in *Chk2*^*-/-*^ cells by treating the cells with etoposide, a topoisomerase II inhibitor which can cause DNA damage. The data showed that, compared to the MEF from wildtype mice, *Chk2*^*-/-*^ MEF cells generated about 5-fold higher *Ifnb* expression (**Fig**. S4A), suggesting that CHK2 could negatively regulate interferon production from the damaged DNA. Consistently, when *Chk2*^*-/-*^ bone marrow-derived macrophage (BMDM) was stimulated by an exogenous dsDNA reagent HT-DNA that could mimic the endogenously damaged DNA fragment, it produced more *Ifnb* gene transcripts and expressed more IFNB protein (**Fig**. S4B-C). Furthermore, *Il6* expression was also elevated in HT-DNA stimulated *Chk2*^*-/-*^ cells but not in polyI:C stimulated group (**Fig**. S4D-E), suggesting that the mechanism of CHK2-related innate immune response regulation is specific to DNA sensing instead of RNA sensing pathway. Importantly, when cells were stimulated with ZIKV, we also observed a consistent elevation of *Ifnb, Mx1* and *Il6* in *Chk2*^*-/-*^ MEF. (**Fig**. S5). Consistent with *in vitro* data, the expression of innate immune genes like *Ifnb, Mx1* and *Il6* in *Chk2*^*-/-*^ mice was significantly elevated as well compared to the littermate control mice upon ZIKV infection (**Fig**. 5E-G).

To further understand if the enhanced innate immune response to ZIKV infection in *Chk2*^*-/-*^ cells was mediated by STING, a sensor of damaged DNA in the cytosol, we infected MEF from wild type, *Chk2*^*-/-*^ and *Chk2*^*-/-*^*Sting*^*-/-*^ mice with ZIKV in parallel. Surprisingly we found that the expression level of *Ifnb, Mx1* and *Il6* returned to the same level as those from wildtype MEF (**Fig**. S5). To determine if STING played a role in protecting *Chk2*^*-/-*^ mice from ZIKV induced pathogenesis *in vivo, Chk2*^*-/-*^*Sting*^*-/-*^ double knockout neonatal mice were infected with ZIKV intracranially (**Fig**. 6A). We observed a significant decrease of survival profile for *Chk2*^*-/-*^*Sting*^*-/-*^ mice as compared to *Chk2*^*-/-*^ mice (**Fig**. 6B). Consistently, the viral load in *Chk2*^*-/-*^*Sting*^*-/-*^ mice was bounced back to the same level as that in the wildtype mice (**Fig**. 6C). Collectively, these data indicate that the STING pathway plays an essential role in sensing damaged cytoplastic DNA from ZIKV infected *Chk2*^*-/-*^ mice, generating interferon and proinflammatory cytokine to protect *Chk2*^*-/-*^ mice from ZIKV induced pathogenesis.

**Figure 6.**
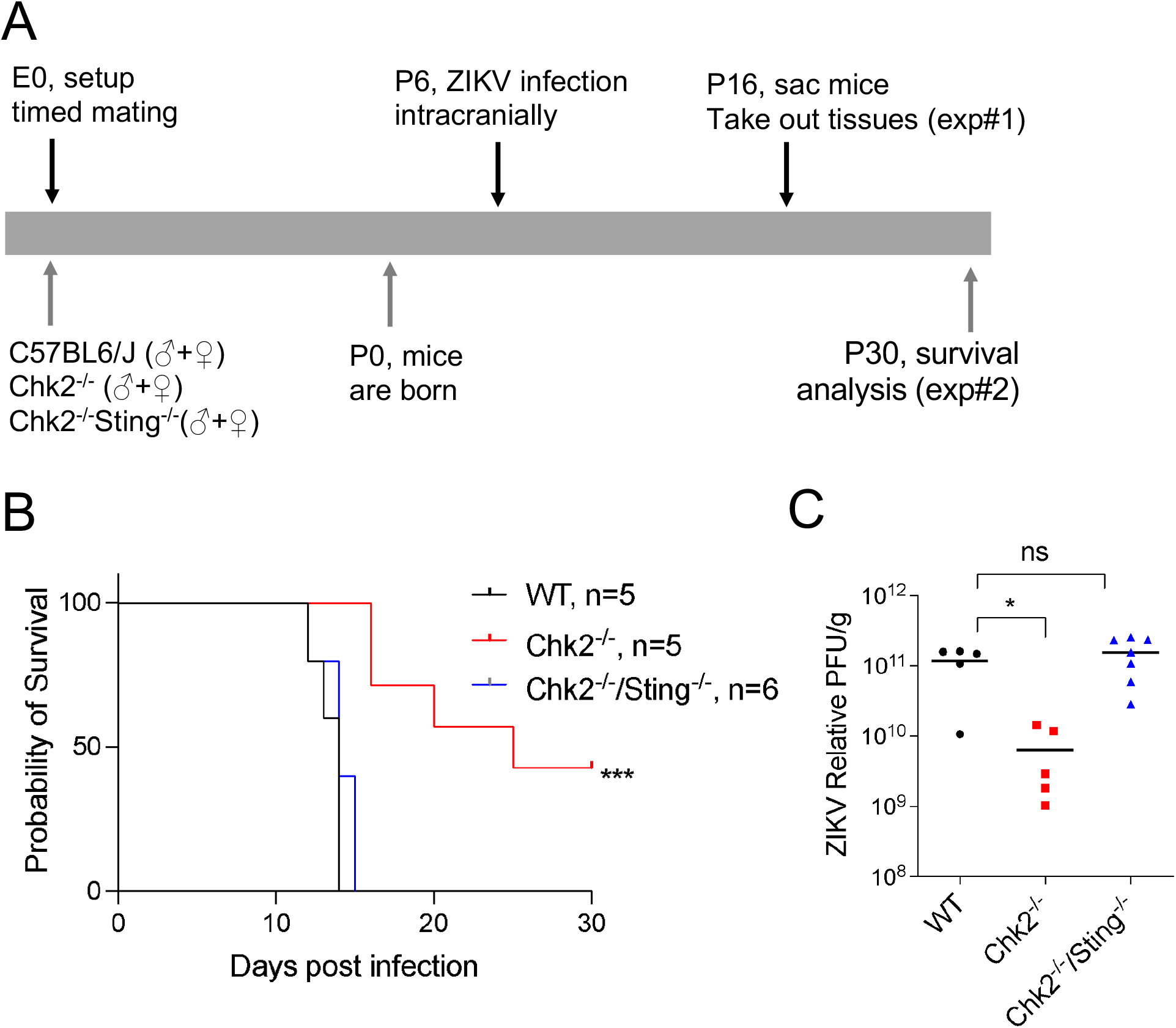
STING pathway is important for enhanced interferon and proinflammatory response after ZIKV infection in *Chk2*^*-/-*^ mice. **(A)** The outline of the experiment. **(B-C)** Survival and viral loads of WT, *Chk2*^*-/-*^, *Chk2*^*-/-*^ *Sting*^*-/-*^ neonatal mice infected by ZIKV (MR766, 100 PFU/mice) were shown. * p<0.05, *** p<0.001, Log-Rank (Mantel-Cox) test (B), or one-way ANOVA test (C).

## Discussion

The results here show that ZIKV infection could uniquely regulate ATM-CHK2-γH2AX DDR pathway to create a beneficial environment for its infection and pathogenesis compared to DENV-2 and YF-17D. When CHK2 is inhibited by small molecule inhibitor or genetic knockout, the beneficial effect is reduced whilst the host antiviral response is elevated through a STING dependent manner. Mechanistically, our data suggest that activation of CHK2 axis by ZIKV would promote the repairment of damaged DNA and prevent its cytoplasmic release to evade STING-mediated antiviral effect.

Regarding the type of DNA damage induced by ZIKV and the related mechanisms, the increased phosphorylation of γH2AX in our *in vitro* and *in vivo* studies suggest it is from DNA DSBs (**Fig**. 1C and **Fig**. 1F). This is also reported by Hammack et al from ZIKV infected hNPCs^16^, revealing that ZIKV induces DSBs consistently in both human and mouse cells across different cell types. Several viruses from other family members are also reported to induce DSBs and regulate cell cycle progression, however the mechanisms involved are different ^20, 21, 22^. γ-herpesviruses 68 (γHV68) and Epstein-Barr virus (EBV) are both DNA viruses, the transient expression of their viral kinases orf36 and BGLF4 alone are sufficient to induce γH2AX, suggesting the viral DNA replication intermediates are not the only sources to induce DDR^23^. Hepatitis C virus (HCV), an RNA virus from Flaviviridae is found to use NS3 protein alone to induce DSBs and enhance mutation frequency of cellular genes^24^. For ZIKV, the related mechanisms of how DSBs are induced are still not clear and identification of the viral or cellular factors controlling DSBs will be of interest to explore further. Regarding the tissue sites where DSBs occur, we observe prominent phosphorylation of γH2AX at both placenta and fetus, the two places that manifest the most obvious ZIKV clinical presentations ^8, 9^. This suggests that inhibition of DDRs at these sites may play a crucial role in controlling ZIKV vertical transmission through placenta and restoring fetal brain developmental defects. Consistently, from our studies in the pregnant mouse infection model, the viral load at placenta is dramatically suppressed by CHK2 inhibitor, and fetus developmental outcomes are improved obviously as evidenced by reduced embryo resorption rate and increased gain of fetal body weight. Furthermore, the viral load in the brain tissues from the intracerebral mouse infection model is decreased dramatically by over 1 log in the *Chk2*^*-/-*^ mice in comparison to the littermate controls. Therefore, targeting CHK2 axis at both placenta and fetal brain through small molecule inhibitors may represent a new approach to control ZIKV infection and the resulted pathogenesis.

Another intriguing aspect of the study is that we reveal a mechanism for how inactivation of CHK2 axis would inhibit ZIKV infection and pathogenesis. We find that when *Chk2*^-/-^ mice are challenged by ZIKV in different infection models, the viral load at different regions of the brain is reduced whilst the antiviral immune response is elevated. Strikingly, when *Sting* is knocked out in the context of *Chk2*^*-/-*^ mice, the antiviral effect is disappeared, and the viral load is bounced back to the same level as that from the wildtype mice. Since CHK2 is one of the major regulators to mediate downstream DNA damage repair, we hypothesized that when the function of CHK2 is blocked, the damaged DNA is unable to be repaired efficiently so that cytoplasmic release of the damaged DNA could trigger STING signaling pathway, resulting in enhanced antiviral innate immunity. Among the three flaviviruses we tested, ZIKV seems to have evolved a unique strategy to evade STING sensing mechanism by activating CHK2 axis to repair the damaged DNA. Another example related to STING activation via CHK2 is in these tumors with deficiency or mutation in ARID1A, which targets CHK2 for ubiquitination and degradation^25^. When CHK2 axis is inhibited in this type of tumor by small molecule inhibitors, the increased DNA replication stress induces DNA damages and release them into cell cytosol to activate STING, promoting lymphocytes infiltration to clear the cancer cells. It is also reported that Japanese encephalitis virus (JEV) induces G1 phase arrest and CHK2 activation, knockdown or inhibiting CHK2 reduces JEV replication^26^. Similarly, HCV induces G2/M phase arrest and activate DDRs, inactivation of ATM or CHK2 suppresses its RNA replication^27^. Whether JEV or HCV uses a similar mechanism to evade STING sensing via CHK2 axis would be of interest to study in this *Chk2*^*-/-*^*Sting-/-* mice model. Since both JEV and ZIKV induce neuropathogenesis, whether CHK2 activation would link to this pathogenic phenomenon will be of great interest to explore too.

In summary, these results reveal some novel mechanistic insights into games that ZIKV plays with the host. ZIKV stimulates DDR, which in turn results in activation of CHK2 and cell cycle arrest, likely to maintain the genomic integrity of the cell. However, the collateral cost to the host of this homeostatic response is evasion of STING-dependent innate immunity by ZIKV, as a result of DNA repair mechanisms orchestrated by CHK2. Thus, ZIKV infection appears to represent an evolutionary détente of sorts: the host cells initiate a DNA repair and cell cycle program to preserve the integrity of the host genome, whilst suffering the lesser calamity of muting the STING antiviral defense mechanism. These results also highlight the pharmacological modulation of the CHK2 pathway as a novel therapeutic strategy against ZIKV.

## Materials and Methods

### Ethics statement

All animal studies were conducted by following animal protocols reviewed and approved by the Stanford University Administrative Panel on Laboratory Animal Care. Mice were bred and maintained in a specific pathogen free Research Animal Facility at Stanford University Department of Comparative Medicine in accordance with guidelines of the US National Institutes of Health. Inoculations of virus were performed under anesthesia to minimize animal suffering.

### Cells, Viruses and other reagents

Hela cells and Vero cells were purchased from ATCC and maintained in DMEM (Thermo Fisher Scientific) containing 10% FBS (Optima, Atlanta Biologics). Zika virus strains PRVABC59 (ZIKV-PR) and MR-766 (MR-1947) were obtained from Dr. Mehul Suthar’s lab ^2^. YF-17D subpassaged from YF-VAX (Aventis Pasteur) in SW-480 cells was a gift from Dr. Ahmed ^28^. DENV serotype 2 virus (strain 16681) was a gift from Dr. Perng as described before ^29, 30^. All virus strains were passaged once in Vero cells cultured in DMEM supplemented with 10% FBS to generate the working viral stocks. Viral stocks were titrated by a standard plaque assay on Vero cells and stored at -80°C as described previously ^2^.

The compounds used in this study were: Etoposide (Sigma-Aldrich, E1383), ATM inhibitor KU-55933 (Sigma-Aldrich, #SML1109), CHK2 inhibitor II (CHK2i) (Sigma-Aldrich, #C3742), CHK2 inhibitor PV1019 (EMD Millipore, #220488).

### Mice and ZIKV infection experiments

C57BL/6 (WT), *Ifnar1*^-/-^ mice were purchased from Jackson Laboratories. *Chk2*^-/-^ mice were kindly provided by Dr. Tak W Mak (University Health Network, Toronto) and were generated as previously described ^18^. *Chk2*^+/-^ breeders were generated by crossing *Chk2*^-/-^ with C57BL/6 (WT) mice. By breeding *Chk2*^+/-^ mice, the obtained *Chk2*^-/-^ mice and littermate controls were used in the studies. In some studies, C57BL/6 mice were used to substitute the littermate controls.

In the pregnant mice infection model, mice were set up for timed mating as described before ^9^. At embryonic days E8.5, the pregnant mice were inoculated by subcutaneous route in the footpad with 1000 pfu of ZIKV-PR strain in 50μl of PBS. Mice were sacrificed at E15.5 to harvest the placentas and fetuses for viral load determination and other assays.

In the neonatal mice infection model, 6-day-old *Chk2*^*-/-*^ mice and littermate controls were injected with 100 PFU ZIKV-PR strain in 5μl of DMEM by intracerebral route. The survival data were recorded daily. On day 10 post infection, mice were sacrificed to harvest the brain for viral load determination and immunofluorescence assay.

In the adult mice infection model, 6-week-old *Chk2*^*-/-*^ mice and littermate controls were injected with 10^4^ pfu ZIKV-MR766 strain in 20μl of DMEM by intracerebral route. The mice were sacrificed on day 10 post infection to harvest the brain for viral load determination and immunofluorescence assay.

### Antibodies and Western blotting

Antibodies against the following proteins were used: Phospho-CHK2 (Thr68) (Cell Signaling, #2197), Total CHK2 (Cell Signaling, #6334S), Phospho-ATM (Ser1981) (Cell Signaling, #5883), Total ATM (Cell Signaling, #92356S), Phospho-Histone H2A.X (Ser139) (Cell Signaling, #9718), Total Histone H2A.X (Cell Signaling, #7631S) and Tubulin (Cell Signaling, #2144). HRP-conjugated anti-mouse (Thermo Fisher Scientific, #G-21040) and anti-rabbit (Thermo Fisher Scientific, #G-21234) antibodies were used as secondary antibodies.

Antibodies against the following proteins were used in Figure S1: Phospho-Rb (Ser807/811) (Cell Signaling, #9308), Cyclin B1 (Cell Signaling, #12231Cyclin D3 (Cell Signaling, #2936), Cyclin E2 (Cell Signaling, #4132), CDK2 (Cell Signaling, #2546), CDK4 (Cell Signaling, #12790), CDK6 (Cell Signaling, #3136), CDK1 (Phospho-CDK1, Cell Signaling, #4539), CDC25C (Cell Signaling, # 4688), Phospho-CDC25C (Thr48) (Cell Signaling, #9527) and Phospho-Histone H3 (Thr11) (Cell Signaling, #9767).

For immunoblotting, cells or tissues were collected in ice-cold RIPA lysis buffer (Cell Signaling, #9806) supplemented with 1x protease/phosphatase inhibitor cocktail (Cell Signaling, #5872). Samples were homogenized with ceramic beads via BEAD RUPTOR 4 (Omni International). After centrifugation, the supernatants were collected for protein quantification via BCA assay (Pierce™ BCA protein assay kit, #23225). Equal amounts of protein from whole lysates were run on an SDS-PAGE and transferred onto nitrocellulose membranes. After blocking with 5% fat-free milk, the membranes were incubated at 4°C with the indicative primary antibodies and secondary antibodies. Proteins were visualized with SuperSignal™ West Femto substrate (Thermo Fisher Scientific, #34095), and the signals were acquired and analyzed via Odyssey Fc (LI-COR) or Chemi Doc MP (Bio-Rad) systems.

### Immunofluorescence assay

Cells were fixed with 4% paraformaldehyde (PFA) for 10 minutes, then permeabilized in PBS containing 0.5% Triton X-100 for 10 minutes. After blocking with 1% BSA in 0.5% Triton X-100 for 30 minutes, the cells were stained for 1 hour with antibody against flavivirus E protein (Novus, clone 4G2, #NBP2-52709) and G3BP1 (Proteintech, #13057-2-AP), respectively. Samples were then stained with Alexa Fluor 488-conjugated goat anti-mouse (Thermo Fisher Scientific, #A-11001) or Alexa Fluor 555-conjugated Donkey anti-rabbit (Thermo Fisher Scientific, #A-31572) secondary antibodies. Nuclei were stained with 4’,6-diamidino-2-phenylindole (DAPI) (Thermo Fisher Scientific, #D1306), and images were captured with LSM 880 confocal (Carl Zeiss) or BZ-X810 (KEYENCE) fluorescence microscope.

### Immunohistochemistry

Placenta or brain tissues were fixed in 4% PFA, then dehydrated in 30% sucrose, and embedded in optimal cutting temperature (OCT) medium, then were frozen and cut into thick sections. Immunofluorescence staining was performed on sections (thickness: brain, 40μm; placenta, 10μm) as described previously^31^. The antibodies used for immunostaining were anti-ZIKV (Genetex, #GTX634157), activated-caspase3 (Abcam, #ab2302). DAPI was used to stain the nuclei.

### Cell cycle analysis and flow cytometry

To precisely profile cell cycle at each phase, the cells were analyzed by bromodeoxyuridine (BrdU) incorporation and DNA content staining with APC BrdU Flow Kit (BD Biosciences, #552598) per the manufacturer’s instructions. Briefly, the cells were infected with the indicated viruses at MOI of 0.1 for 36 hours. Immediately prior to collection, cells were incubated with 10 µM BrdU for one hour, followed by washing with PBS. Afterwards, the cells were stained with LIVE/DEAD™ Aqua fixable dye (Thermo Fisher Scientific, #L34957) for 30 minutes to distinguish live/dead cells. Then the cells were fixed with BD Cytofix/Cytoperm buffer for 30 minutes, and permeabilized with BD Cytoperm Permeabilization buffer for 10 minutes. After washing, an additional 5 minutes re-fixation with BD Cytofix/Cytoperm buffer was applied on ice for 5 minutes. To expose the incorporated BrdU for antibody staining, the cells were then treated with 300 μg/ml DNase for one hour at 37°C. Afterwards, BrdU was stained with APC-conjugated anti-BrdU antibody for 20 minutes at room temperature. After additional washes, the cells were resuspended in 7-amino-actinomycin D (7-AAD) solution, followed by data acquisition on a BD LSR II flow cytometer at a rate no greater than 400 events per second. The data were analyzed using FlowJo software.

### Virus titration assay

The titer for ZIKV-PR, DENV-2 and YF-17D was determined by plaque assay on Vero cells. Basically, Vero cells were plated in 24-well plates the day before infection. On the infection day, the virus samples were 10-fold serial diluted in culture medium, then 250uL of sample from each dilution was inoculated in the pre-washed Vero cells. After 2 hours incubation at 37°C, the unbound viruses were wash away by cell culture medium. Then 0.5% agarose gel containing 5% FBS in DMEM was added on top the cell monolayer. After 3 days incubation at 37°C, remove the plate and add 2mL 4% PFA to each well to fix the cells for 1 hours. Afterwards, both PFA and agarose were removed and then stained with crystal violet followed by wash with water. The number of plaques were counted and used for virus titer calculation.

### Quantification of viral load

Total RNA from tissues, cell lysis or cell supernatants was extracted with the PureLink™ RNA Mini Kit (Thermo Fisher Scientific, #12183052) per the manufacturer’s instructions. For viral load detection, quantitative reverse transcription PCR (qRT-PCR) was conducted using Luna universal probe one-step RT-PCR kit (NEB, #E3006) on CFX96 Touch Real-Time Detection System on 96-well plates (Bio-Rad). The specific primers and probes targeting the conserved region of viral NS gene were designed and available upon request. Viral relative PFU was calculated based on the standard curve of ct from the serial diluted standard samples with known PFU titer. Viral relative PFU was normalized with tissue weight and plotted with Prism GraphPad software.

### Statistical analysis

All results are displayed as mean ± SEM. Biological replicates were used in all experiments unless stated otherwise. Statistical significance was determined by unpaired student t-test, one-way ANOVA, or two-way ANOVA using Prism software (GraphPad) depending on the experimental layout. Survival analysis was determined by Log-rank (Mantel-Cox) test. Probability values of p < 0.05 were considered significant and denoted by *. Where indicated, ** denotes p < 0.01, *** denotes p < 0.001 and **** denotes p < 0.0001.

## Acknowledgement

The study was supported by NIH grants (R37 DK057665, R37 AI048638, U19 AI090023, U19 AI057266 (PI. Rafi Ahmed), U19 AI159840 (PI, Steven Foung)), the Bill and Melinda Gates Foundation, and the Soffer Fund endowment and Open Philanthropy to BP. We thank members of the Pulendran laboratory for discussion. We also thank the Stanford Shared FACS facility for providing access to the equipment.

## Author Contributions

H.L.M. designed and executed experiments, wrote the first draft of the manuscript. C.F.L. designed and executed experiments, prepared the manuscript. Y.L.W., E.V.V., P.H., and J.R.B. collaborated for experiments execution. M.T., and H.S.H. genotyped and took care of the mice. T.W.M. and M.S.S. provided reagents and scientific collaboration. B.P. conceived the whole study, guided experimental design and wrote subsequent drafts of the manuscript.

## Supplemental Figures

**Figure S1.**
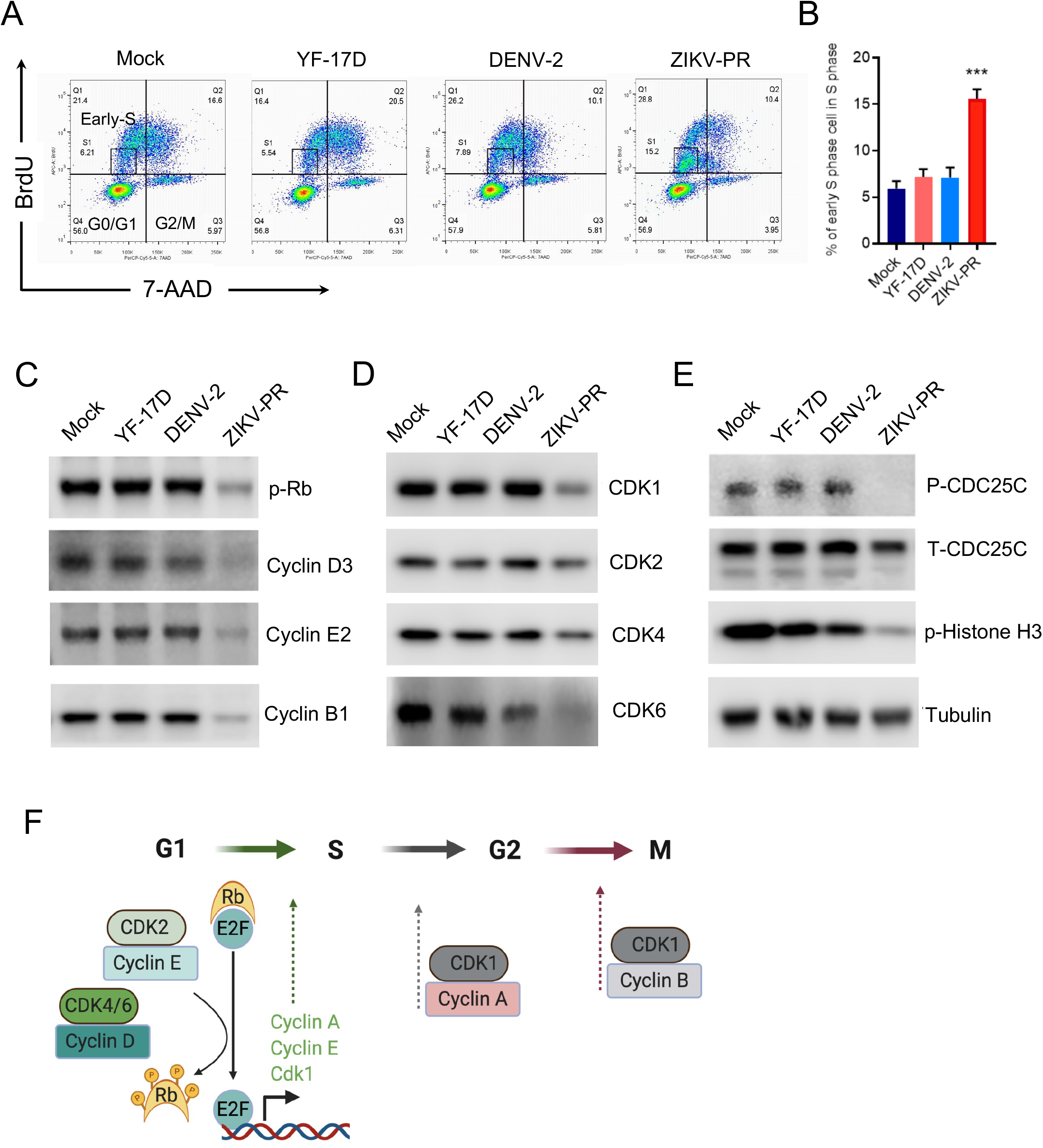
ZIKV arrests cell cycle at early-S phase by impairing multiple key cell cycle regulators. **(A-B)** Hela cells were infected with ZIKV-PR, DENV-2 and YF-17D viruses at MOI of 0.1 for 36 hours. The cells were then pulsed with BrdU for 1 hour and followed by co-staining with 7-AAD. The cell cycle profile and the percentage of cells at early-S phase were represented from 2 independent experiments. Error bars are mean ± SEM, average of three biological replicates, *** p<0.001. **(C-E)** Hela cell lysates from the aforementioned experiments, but without BrdU pulse were subjected to western blot analysis of the key regulators that control cell cycle progression. One set of representative images from two independent experiments were shown. **(F)** Cartoon of checkpoint regulators involved in each step of cell cycle process.

**Figure S2.**
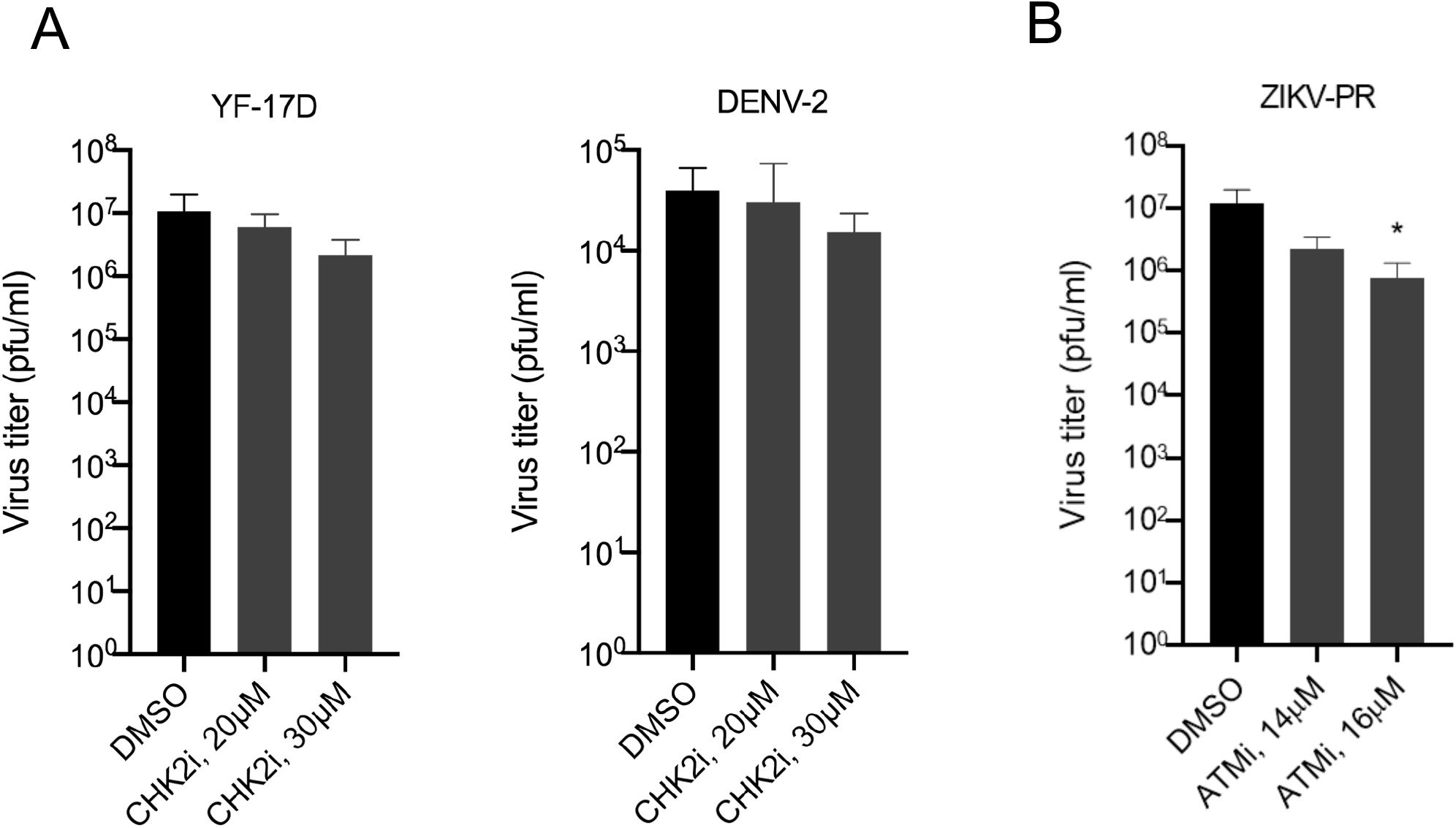
The effect of CHK2 and ATM inhibitor on virus infection *in vitro*. **(A)** Hela cells were infected with DENV-2 or YF-17D virus at MOI of 0.1, then treated with two different doses of CHK2 inhibitor. The supernatants were collected at 48 hours post infection for plaque assay. **(B)** To determine the effect of ATM inhibitor on ZIKV infection, Hela cells were infected with ZIKV at MOI of 0.1, then treated with ATM inhibitor at different doses that do not induce cytotoxicity. The supernatants were collected at 48 hours post infection for plaque assay. The plotted data were pooled from 2 independent experiments. Error bars are mean ± SEM, * p<0.05, one-way ANOVA test.

**Figure S3.**
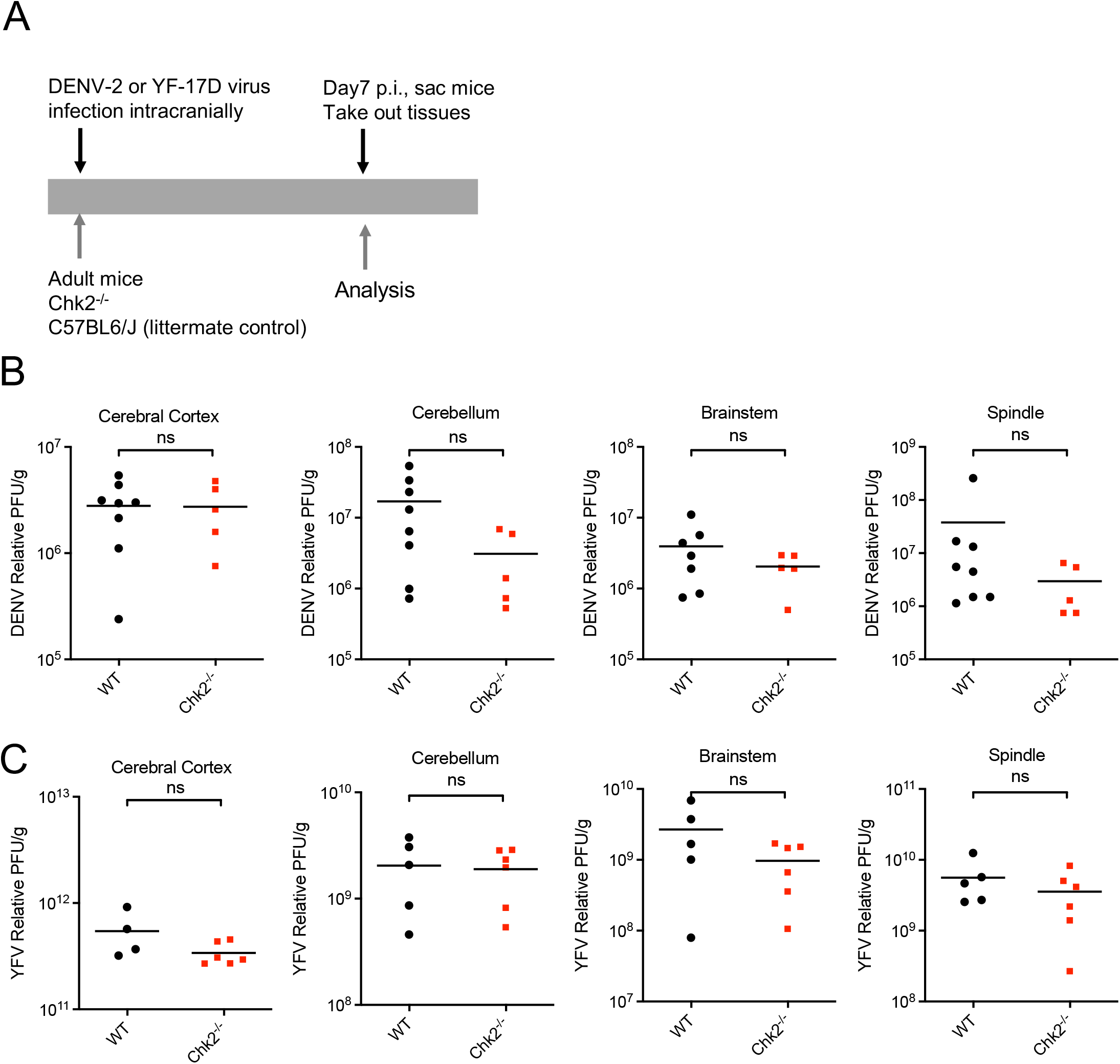
The effect of *Chk2* knockout on Dengue virus and yellow fever virus infection *in vivo*. **(A)** Schematic of virus infection in the brain of adult mice. **(B-C)** 6-week-old adult *Chk2*^*-/-*^mice or littermate controls were infected with DENV-2 (**B**) or YF-17D (**C**) at 10^4^ PFU/mouse by intracranial injection. At 7 days post infection, the viral loads in the cerebral cortex, brainstem, cerebellum and spinal cord were detected by qRT-PCR, ns: non-significant, student t test.

**Figure S4.**
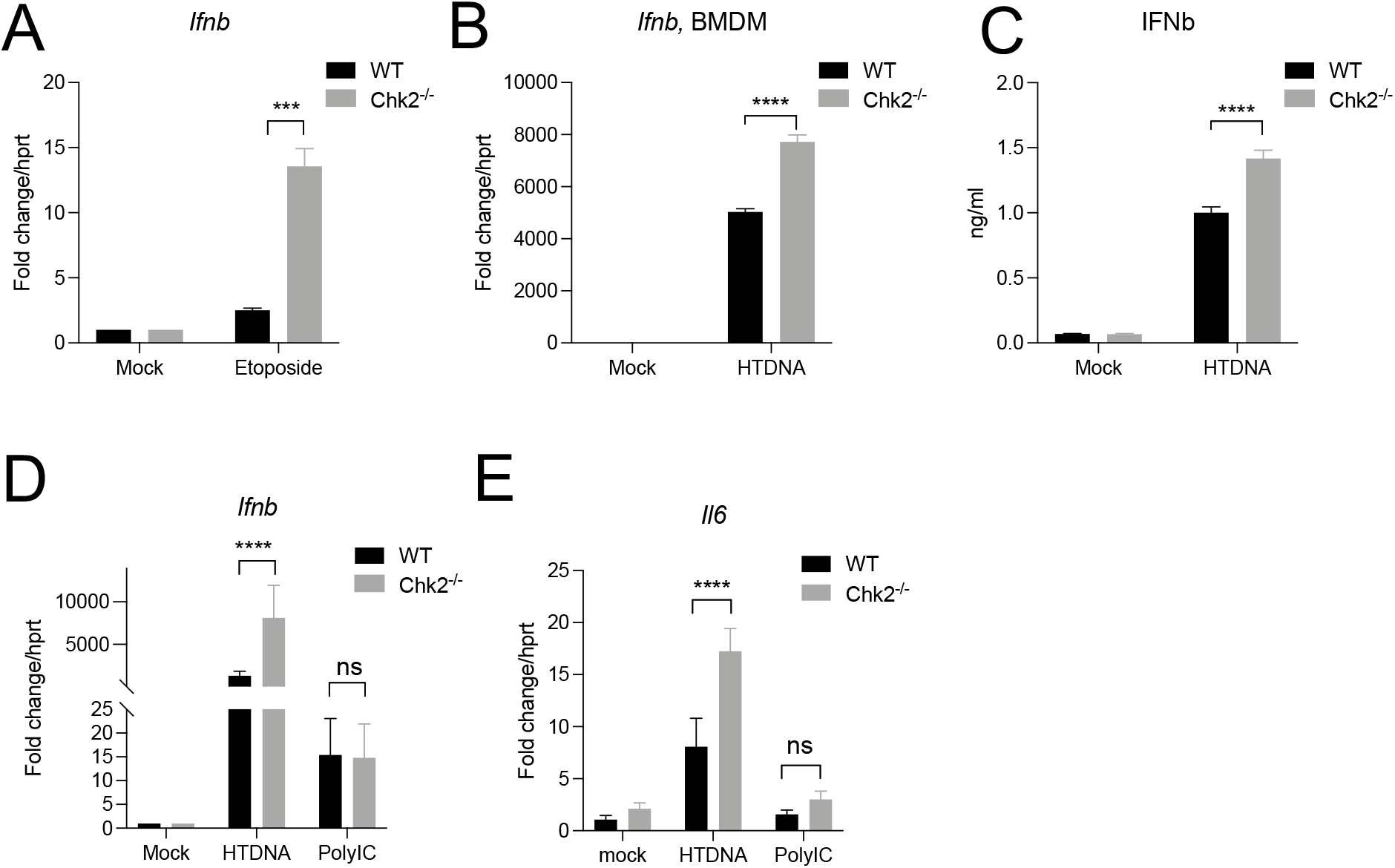
Immune response in *Chk2*^*-/-*^ cells stimulated by Etoposide, DNA or RNA. **(A)** Interferon response and ISG production in *Chk2*^*-/-*^ cells treated by Etoposide. **(B-D)** The Ifnb expression in WT and *Chk2*^*-/-*^ cells stimulated by DNA or RNA ligands at 12 hours post stimulation. **(E)** The Il6 expression in WT and *Chk2*^*-/-*^ cells stimulated by DNA or RNA ligands at 12 hours post stimulation. Representative data was shown from two repeats, *** p<0.001, and **** p<0.0001, one-way ANOVA test (A-E).

**Figure S5.**
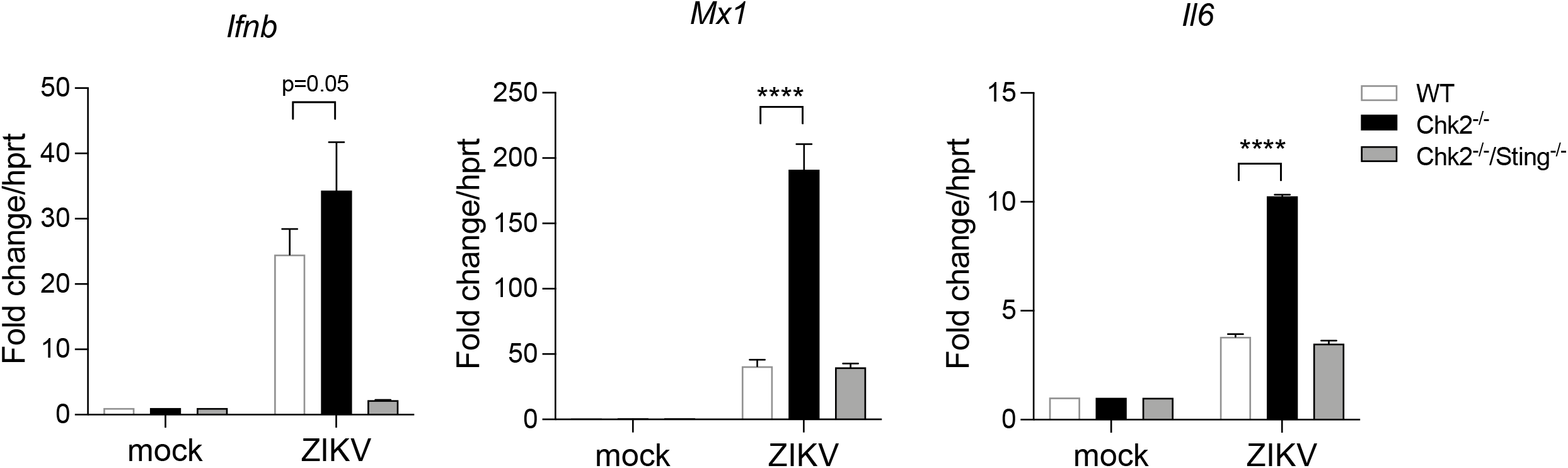
Immune response in *Chk2*^*-/-*^ and *Chk2*^*-/-*^*Sting*^*-/-*^ cells after ZIKV infection. *Ifnb, MX1*, and *Il6* gene expression in wildtype, *Chk2*^*-/-*^, *Chk2*^*-/-*^*Sting*^*-/-*^ MEF cells at 48 hrs after ZIKV infection (MOI=1). Representative data was shown from two repeats, **** p<0.0001, one-way ANOVA test.

